# The relationship of genetic diversity and inbreeding to extinction risk across over 500 vertebrate diploid genomes

**DOI:** 10.64898/2026.08.14.744813

**Authors:** Amanda Gardiner, Vertebrate Genomes Project Phase 1 Consortium, Richard Durbin

## Abstract

Genetics may help address the biodiversity crisis by providing information about genetic diversity and temporal changes in demography for species of interest. Advances in whole-genome sequencing create new opportunities for demographic analysis, even based on the two copies of a genome found in a single diploid individual. The Vertebrate Genomes Project (VGP) is generating high-quality, chromosome-level reference genomes across the full range of extant vertebrate species, with its first phase delivering assemblies spanning approximately 95% of vertebrate orders. Using 512 diploid VGP genomes, we quantified intra-species heterozygosity, runs of homozygosity (ROH), and inferred past effective population sizes (***N_e_***) with the Pairwise Sequentially Markovian Coalescent (PSMC). Threatened species are more likely to exhibit lower heterozygosity and longer ROH, though there is large variation in both measures across all IUCN categories. Interestingly, PSMC suggests that estimated historical ***N_e_*** several thousand generations ago is a better predictor of threatened status than the present day estimate. Co-analysing with life history traits, we found that marine species tend to have lower ROH content, while fossorial species show significantly higher inbreeding levels. Indeed, habitat and foraging strata are much stronger predictors of IUCN status than genetics, with estimated historical ***N_e_*** providing a small but significant amount of additional information. Together, these results suggest that, while measures of genetic diversity are correlated with IUCN status, much of that correlation may derive from ecological factors such as habitat, with only a relatively small direct contribution. Nevertheless, reference genomes like those generated by the VGP can yield valuable information, like historical ***N_e_***, while facilitating population monitoring and management for species of interest.

## 1 Introduction

We are currently in the midst of a biodiversity crisis, with over 28% of all assessed species of plants and animals threatened with extinction (IPBES 2019; IUCN 2025). Conservation genetics (CG) can help address this crisis by identifying genetic threats; such as loss of effective population size (*N_e_*), inbreeding (the mating of related individuals), changes in gene flow between populations, and expression of recessive deleterious alleles; with the information feeding into decisions about genetic management (Weeks et al. 2017; Willi et al. 2022; Kardos et al. 2021; Theissinger et al. 2023; Speak et al. 2024; Zheng et al. 2026).

Two common factors studied in CG are genetic variation and inbreeding. Low genetic variation in a population is associated with bottlenecks and/or long-term low effective population size, arguably increasing extinction risk, while high variation is said to be associated with higher fitness and decreased risk due to higher levels of potential adaptability (Reed and Frankham 2003; Theissinger et al. 2023; Willi et al. 2022; Kardos et al. 2021; Willoughby et al. 2015; Leigh et al. 2019; Romiguier et al. 2014). The standard measure of genetic diversity is population heterozygosity, but there is debate about how strong the correlation really is between heterozygosity and extinction risk (Teixeira and Huber 2021; Brüniche-Olsen et al. 2018, 2019; Doyle et al. 2015; Grossen et al. 2020). Inbreeding can lead to inbreeding depression, which is the loss of reproductive fitness in a species due to expression of deleterious recessive alleles. Inbreeding can be estimated through looking at runs of homozygosity (ROH), which are long stretches of homozygous genotypes in an individual’s genome assumed to be autozygous in origin and inherited from a relatively recent common ancestor (Ceballos et al. 2018). The longer ROH are, the more recent the common ancestor between parents is, as ROH get broken up with every successive generation due to recombination (Ceballos et al. 2018). Early inbreeding research primarily focused on quantifying ROH in humans and livestock (for examples see Tao et al. (2025); Pegolo et al. (2025); Marras et al. (2015); McQuillan et al. (2008); Pemberton et al. (2012); Szpiech et al. (2013); Nothnagel et al. (2010)). Recently similar approaches have been used to quantify inbreeding and genetic load in endangered species (Kardos et al. 2023; Furni et al. 2025; de Greef et al. 2024; Macciotta et al. 2021; Kyriazis et al. 2025b; Hewett et al. 2023a; Stanhope et al. 2023)).

Detection and analysis of ROH, however, requires comparison of high-quality chromosomal scale genetic data. Until recently, this was a limiting factor in CG research due to the complexity and cost of sequencing non-model species (Li and Durbin 2024; Cheng et al. 2024). This situation is however changing rapidly, with efforts globally coordinated for all eukaryotes by the Earth Biogenome Project (Blaxter et al. 2025), and for vertebrates by the Vertebrate Genomes Project (VGP). The VGP is a worldwide scientific collaboration launched in 2017 which aims to create high-quality chromosome-level reference genomes for the over 70,000 extant vertebrate species on earth (Rhie et al. 2021), using modern long-read genome assembly approaches that provide the sequences of both chromosome sets in a diploid organism (Li and Durbin 2024). The project has now completed its first phase, covering over 85% of vertebrate orders in 579 species (Formenti et al. 2026). Initial results using data from 219 of these species suggest that as a species becomes more threatened they tend to have higher median levels of ROH and lower median heterozygosity (Formenti et al. 2026).

Here we expand upon those initial results to further understand how genetic diversity and ROH are related to extinction risk. We extended the dataset to include 512 VGP species by incorporating additional data, ran MSMC2 (Schiffels and Durbin 2014) to understand how *N_e_* relates to genetic diversity, quantified the patterns of short and long ROH to see whether species with similar extinction risk have similar demographic histories, and looked at life history traits to understand how those correlate with diversity and ROH. These results provide a broad picture across vertebrates of the relationships between genetic diversity and IUCN status; in some cases providing information that may be relevant for genetic management of specific species, while more broadly providing analysis tools and background data for conservation genetics and population management.

## 2 Materials and Methods

### 2.1 Included Species

The first VGP data freeze contains 579 primary genomes, representing over 85% of vertebrate orders across all major vertebrate lineages. For details on species, sampling, and sequencing see (Formenti et al. 2026). All assemblies are diploid, with the secondary assemblies varying in completeness depending on when it was made and the methodology used for assembly (Formenti et al. 2026). For species where the secondary assembly did not meet VGP criteria (*>* 1Mb contig N50, BUSCO gene completeness *>* 95%), we used the VGP secondaries if they met completeness criteria of *>* 90% coverage. For species where this was not the case, we manually went through all publicly available assemblies for a given species and picked the best quality assembly as a secondary based on assembly level, contig N50, total assembly length, and coverage of the primary. In total, there are 89 additional secondary genomes, and therefore in these species the primary and secondary genomes come from different individuals. The vast majority of these additional genomes were sequenced either as part of the VGP or at the Wellcome Sanger Institute as part of the Darwin Tree of Life Project. A list of accession numbers for all genomes used is available in the Supplementary Data.

#### 2.1.1 Life History and Captivity Data

A diploid genome only provides a random sample of wild population genetic data if the individual was bred in the wild. Multiple genomes in the VGP were sampled from individuals in captivity, however in many cases the genome metadata does not specify whether the individual was captured from the wild or bred in captivity. Therefore, any genome was flagged as “captive” if it came from a captive-living or domesticated individual. Captivity data for the individuals used to generate the VGP assemblies was provided by Peter Sudmant and Runyang Nicolas Lou (Formenti et al. 2026). Sampled individuals were determined to be in captivity if that was either clearly stated in the metadata or the sampling location was outside the range for the species. We collected captivity data for non-VGP samples from the metadata associated with the assembly at NCBI, based on provided information, location (whether it was sampled outside the species range), and associated publications. The data, with captivity flags and notes justifying captivity flag, are provided in the Supplementary Data. If the primary and alternate assemblies were from different individuals and had different captivity status, the status for the primary assembly was used to filter the data.

IUCN Red List status and habitat information were obtained using the IUCN redlist package *rredlist*, with manual verification of IUCN status (Gearty et al. 2025). All other life history information including stratum was obtained either through FishBase for all cartilaginous and bony fish, or using the TetrapodTrait database for all other species (Moura et al. 2024; Boettiger et al. 2012).

### 2.2 Genome Alignment and Variant Calling

Genome alignment was done using *FastGA*, followed by chaining to find maximal colinear subsets of alignments with *ALNchain*, and converting the output pairwise alignment file to a PAF file (Myers et al. 2024). Genome alignment and subsequent downstream analyses were all done using a custom Snakemake pipeline (Mölder et al. 2021). After alignment, we removed all alignments which had abnormally high divergence (*≥*0.1), and/or mapped onto unplaced scaffolds. *Paftools* was used to find uniquely aligned regions of the genome, stored as a BED file, and heterozygous sites, stored in a VCF file (Li 2018). The resulting VCF was filtered to remove indels and structural variants, only retaining single nucleotide variants (SNVs).

### 2.3 Runs of Homozygosity

ROH were detected from BED and VCF files using a custom script. On a given chromosome we measured the distance between variants in bases, only counting bases that were aligned:

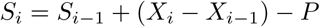

where *S_i_*is the length ‘score’ for ROH at any given variant, *S_i−_*_1_ is the score at the previous variant, (*X_i_ − X_i−_*_1_) is the distance in aligned bases between the current variant and previous variant, and *P* is a penalty threshold that must be passed to be considered an ROH, which we set as 100,000. Therefore ROH segments are defined as being a minimum length of 100 kilobases (kb). Equivalently, SNVs more than 100kb apart are permitted within a called ROH. We calculated the score for each successive variant along a chromosome, with a ROH starting when the score became positive. When variants are more frequent and closely distributed such that the score became zero or negative, that marks the endpoint of a ROH. If there was a gap outside of a ROH, it was not be considered. If a gap was encountered between variants within a ROH, the gap was labelled as within the ROH, but its size was not included as part of the ROH length, which was calculated with aligned bases.

Instead of using genetic map units, which many of our species do not have available, or bases, which can bias calculations due to variation in recombination rate, we defined ROH length by the proportion of the chromosome they are on (boundaries: 0.5%, 1%, 2%, 5%, 10%, 20%, 50%, 100%). Any ROH that were at least 1% of the length of the chromosome they were found on were defined as long ROH, given that is approximately what a 1Mb ROH translates to in human chromosomes.

Two metrics were calculated: number of ROH (*N_ROH_*), and the inbreeding coefficient (*F_ROH_*). *N_ROH_* is the count number of ROH present in a sequence of interest (Ceballos et al. 2018). *F_ROH_* is the proportion of the autosomal genome contained in ROH:

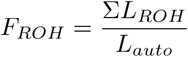

where Σ*L_ROH_* represents the sum of the length (in bases) of all ROH over a defined size across the whole genome, and *L_auto_* represents the length of the aligned autosomal genome in bases (McQuillan et al. 2008). Both ROH calculated and the length of the genome were measured in aligned bases only.

Bayesian regression with phylogenetic correction using *brms* with *Stan* in addition to the phylogeny for the VGP phase 1 data, was done to test for differences in ROH between ordinal IUCN categories, ranked from Least Concern (LC) as one to Critically Endangered (CR) as five (Formenti et al. 2026; Bürkner 2018; Carpenter et al. 2017).

#### 2.3.1 Quantifying ROH patterns

Following Ceballos et al. (2018), we compared *N*_ROH_, which we normalized by the number of autosomal chromosomes, against *F*_ROH_. Typically *N*_ROH_ is compared against the sum length of ROH, however we used *F*_ROH_ as it normalizes across different genome sizes. We compared all individuals with a total *F*_ROH_ of at least 1% across all ROH, and ran a linear regression using the R package *vegan* between the different IUCN categories, testing for statistical significance of IUCN status in comparison to a null model using an ANOVA test (Oksanen et al. 2026).

### 2.4 Heterozygosity

Heterozygosity was calculated in 1Mb sliding windows, with the windows overlapping by 500kb. If a chromosome was shorter than 1Mb, the windows were reduced to 100kb, overlapping by 50kb. If any chromosomes or chromosome fragments were shorter than 100kb, they were excluded from the analysis. The number of variants between the primary and alternate haplotype in each window was summed, excluding ROH:

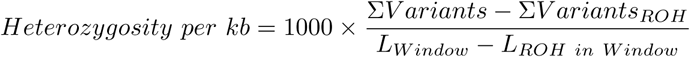

The mean heterozygosity across the whole genome was calculated as the mean of the heterozygosity across all windows on all autosomes.

Bayesian regression with phylogenetic correction using was done to test for differences in heterozygosity outside of ROH between ordinal IUCN categories, ranked from Least Concern (LC) as one to Critically Endangered (CR) as five (Formenti et al. 2026; Bürkner 2018; Carpenter et al. 2017).

### 2.5 MSMC analysis

Multiple Sequentially Markovian Coalescent (MSMC) is a population genetics method which can take an input of phased whole genome sequences and estimate an effective population size for the species for the past several hundred thousand years, up to a few million years ago (Schiffels and Durbin 2014). MSMC2 is an updated software for running MSMC, which can use either a single diploid genome sequence or multiple phased genomes (Malaspinas et al. 2016; Schiffels and Wang 2020). To prepare the data for MSMC2, positive mask files were generated based on the BED files to include only aligned regions, and negative mask files based on the ROH located in the genome so that they were excluded.

The Spearman rank correlation coefficient was calculated for *N_e_*in relation to IUCN status for each time bin. Bayesian regression with phylogenetic correction was done as above to test for correlation between *N_e_* and ordinal IUCN status, for the recent past (approximately 1k generations ago) and two historical time points when the correlation is higher (approximately three thousand generations ago and twenty thousand generations ago).

### 2.6 Analysing the inbreeding coefficient by habitat

Habitat information was extracted from the major habitats in the IUCN habitats classification scheme version 3.1 (https://www.iucnredlist.org/resources/habitat-classification-scheme) using *rredlist* (Gearty et al. 2025). The categories are not mutually exclusive, and if a species is reported in multiple habitats, we scored that species for all relevant habitats. Species were defined as marine if they occurred in at least one of habitats 9-12 (Marine Neritic, Marine Oceanic, Marine Deep Ocean Floor, and Marine Intertidal). We tested for significant differences in long ROH content for marine species while also doing phylogenetic correction using Phylolm and MCMCglmm (Hadfield 2010; Tung Ho and Ańe 2014; Formenti et al. 2026).

Foraging stratum information, defined as ‘categories of habitat use commonly reported’, was obtained from the TetrapodTraits database (TTDB) microhabitat field, and comprises five categories: aerial, arboreal, aquatic, fossorial, and terrestrial (Moura et al. 2024). The categories are not mutually exclusive, and we scored species for all relevant strata. We assigned all non-tetrapod vertebrates, including bony and cartilaginous fishes, in our dataset as aquatic. A Kruskal-Wallis test, followed by a Dunn’s test for subsequent pairwise comparisons, was used to check for significant differences in long ROH content between the different strata (Kruskal and Wallis 1952; Dunn 1964), using R packages *dunn.test*, *MASS*, *vegan*, and *rstatix* (Team 2018; Oksanen et al. 2026; Venables and Ripley 2002; Kassambara 2023; Dinno 2026).

### 2.7 Linear Regression

We created a linear regression model using python package *scikit-learn* to test how well the inbreeding coefficient in a species could be predicted based on other genetic and life-history factors (Pedregosa et al. 2011). A linear mixed effects model was used with the VGP phase 1 phylogeny to account for phylogenetic relatedness. Only species with an *F*_ROH_ value of at least 1%, for all ROH *≥* 100kb were used. The dependent variables in the model were IUCN status (encoded 0 to 4), heterozygosity outside of ROH, historic effective population size from the time bin inclusive of 3,000 generations ago, and habitats and strata (encoded categorically). A similar model for heterozygosity was run with dependent variables IUCN status, *F*_ROH_, habitats and strata. These models were also run only with habitats and strata as coefficients to understand the contribution of genetic factors and IUCN status to the predictions.

A phylogenetically corrected linear regression model was run to predict IUCN status as an ordinal category from Least Concern (0) to Critically Endangered (4) with habitat, strata, *F*_ROH_, heterozygosity outside of ROH, and historical effective population size. The model was then rerun with only habitat and strata as coefficients.

### 2.8 IDRisk

IDRisk is a new metric proposed by (Kyriazis et al. 2025a) to evaluate how susceptible a population will be to inbreeding depression if their population were to decline. It is simply calculated as the product of the inbreeding coefficient for long ROH in a species and the heterozygosity outside of the ROH:

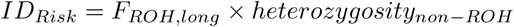

We applied this to our vertebrate data set using the mean heterozygosity per kb outside of ROH regions multiplied by the inbreeding coefficient for ROH that were at least 1% of their chromosome in length. IDRisk is a continuous value, however we also categorised it from low to extremely high risk following the cutoffs set by (Kyriazis et al. 2025a). Values below 0.05 were categorised as low, moderate was defined as being between 0.05 and 0.25, high was set as being from 0.25 to 0.5, while extreme was anything with an IDRisk value of above 0.5.

## 3 Results

We included 512 species in the analysis, all with a primary reference genome sequenced as part of the Vertebrate Genomes Project (VGP), supplemented either by the secondary haplotype from the VGP assembly or by other high quality assemblies (Supp. Table 1). They represented all major vertebrate lineages, with the majority either ray-finned fishes (29%) or mammals (26%) (Supp. Table 2). To evaluate extinction risk, we assigned IUCN Red List status to all species where available. All IUCN statuses from Least Concern to Extinct in the Wild are present; however the distribution of species across categories is uneven, with over 60% of samples in the Least Concern category, and the next most speciose category being Data Deficient or Needing Assessment, which we combined into a single category (Supp. Table 3). To avoid captive breeding programs affecting our results, we filtered out 23% of the dataset containing captive-sampled individuals or domesticated species (Supp. Table 4).

**Table 1.** Bayesian regression results for the relationship of IUCN status to MSMC *N_e_* values through time.

| Number of generations ago | Estimate [95% CI] | pd value (%) | ROPE (%) |
| --- | --- | --- | --- |
| 1,000 | -0.19 [-0.38, -0.03] | 99 | 44.2 |
| 3,000 | -0.26 [-0.49, -0.04] | 99.3 | 22.6 |
| 20,000 | -0.51 [-0.81, -0.21] | 99.9 | 0 |

### 3.1 Heterozygosity and Runs of Homozygosity

Mean heterozygosity outside of ROH across species was 2.78/kb, ranging from a high of 17.3/kb in the Canada goose (*Branta canadensis*), to a low of 0.002/kb in the Anegada rock iguana (*Cyclura pinguis*) and Aeolian wall lizard (*Podarcis raffonei*). We find a negative correlation between heterozygosity and IUCN status, with the exception of Endangered species (Fig. 1A). These same results are recovered when we include captive-sampled species, with a slightly lower mean and median heterozygosity in Endangered species (Fig. S1). Bayesian regression shows that this negative correlation between heterozygosity and IUCN status is statistically significant (*β* = −0.34, 95% CI [−0.55, −0.16], pd = 100%, 2.13% in ROPE), independent of phylogenetic relationships (*σ* = 1.85, 95% CI [0.42, 3.53]).

**Fig. 1.**
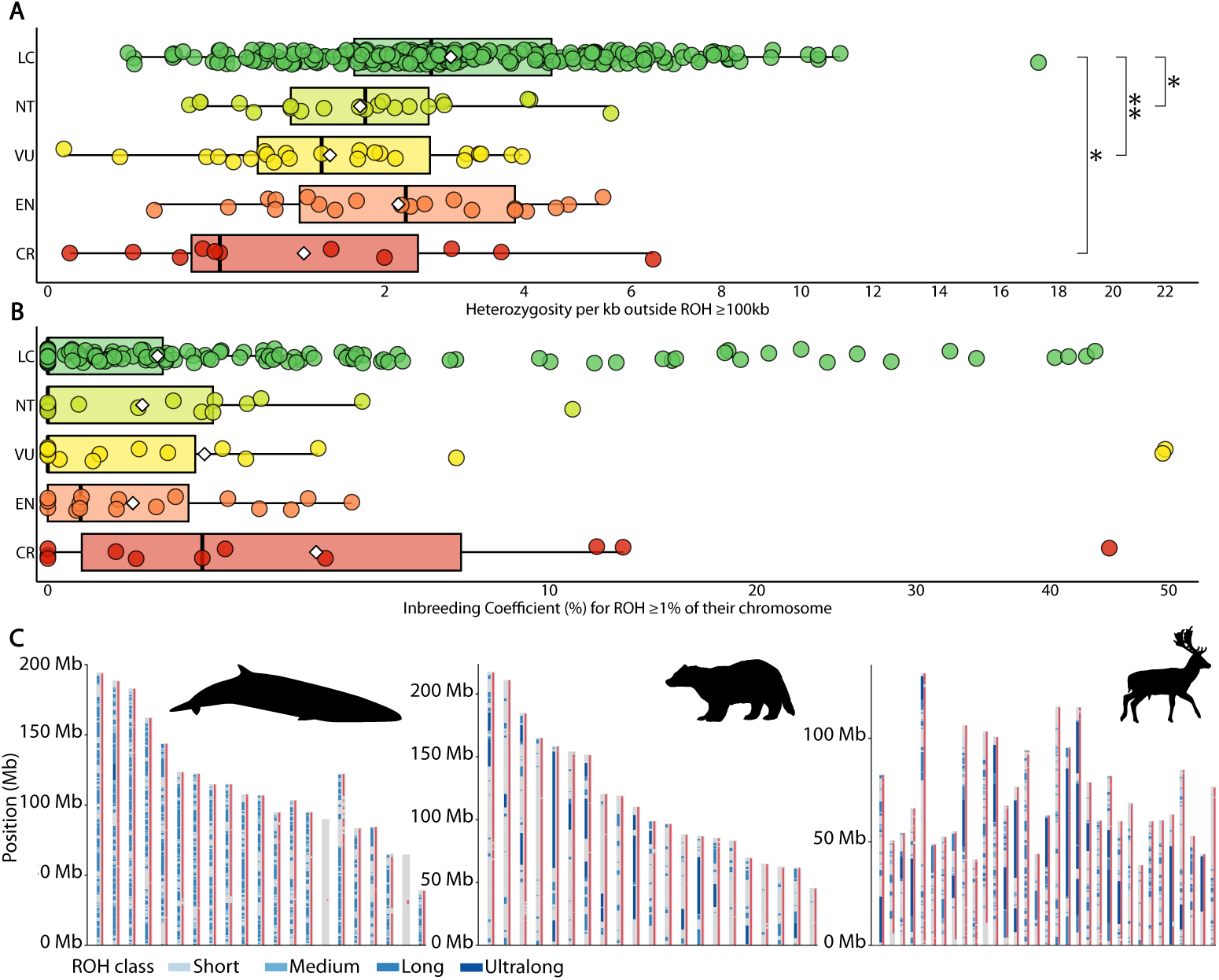
A) Heterozygosity in wild-sampled species. B) The inbreeding coefficient for long ROH, defined as being at least 1% of their chromosome in length, in wild-sampled species. C) Examples of ROH patterns observed in three different mammal species: from left to right *Balaenoptera ricei*, *Meles meles*, and *Dama dama*. The ROH are color coded in different shades of blue based on their length, while the red on each chromosome indicates were the genomes were aligned. LC, Least Concern; NT, Near Threatened; VU, Vulnerable; EN, Endangered; CR, Critically Endangered.

Reciprocally, we find a positive relationship between the inbreeding coefficient (*F*_ROH_) and IUCN status, with *F*_ROH_ for long ROH (defined as being at least 1% of their chromosome in length) increasing as IUCN status increases (Fig. 1B). Bayesian regression showed, however, these differences are not statistically significant (*β* = 0.01, 95% CI [−0.03, 0.05], pd = 73.62%, 100% in ROPE) after accounting for phylogenetic relationships (*σ* = 2.49, 95% CI [1.15, 4.08]). Endangered species have a lower mean *F*_ROH_ than the other IUCN categories, however they have a higher median than all categories but Critically Endangered. No ROH were found in 196 species, while the highest *F*_ROH_ for all lengths of ROH *>* 100kb is in the Suwannee alligator snapping turtle *Macrochelys suwanniensis* (88%), which also had the highest *F*_ROH_ for long ROH (49.8%). The same positive trend between *F*_ROH_ and IUCN status is found when looking at captive-sampled species (Fig. S2). When looking at all ROH (*≥* 100kb) across all species, Endangered species fit into the positive trend, with having the second highest mean and median inbreeding levels behind Critically Endangered species (Fig. S3).

While Least Concern species have the lowest median *F*_ROH_, there are a substantial number of outliers with high *F*_ROH_, showing that other factors besides extinction risk can lead to high levels of inbreeding. Two examples from well-studied species are the European fallow deer (*Dama dama*; 55.8%), and European badger (*Meles meles*; 39.1%), which we compare to the Critically Endangered Rice’s whale (*Balaenoptera ricei*). All three have high levels of inbreeding, but upon closer inspection the ROH patterns between the three species are different (Figure 1C). The fallow deer has evidence of both a past population bottleneck and very recent inbreeding with a bimodal distribution of ROH with both short ROH between 0.17% and 2%, and ultralong ROH that were between 13% and 67% the length of their chromosomes, with a total *F*_ROH_ of 55.8% (Fig. 1C). The badger is indicative of recent inbreeding, with an inbreeding coefficient of 39.1% across all ROH, the majority of which are ultralong ROH between 9% and 100% of the chromosome in length (Fig. 1C). By comparison, Rice’s whale had 73.8% of its genome contained in ROH, with an approximately normal distribution of ROH sizes: the vast majority of ROH are classified from short to long, lying within 0.25% - 4% of their chromosome in length, supporting a small effective population size for an extended period of time (Fig. 1C). These differences between the individuals indicate differences in demographic history between the species both in the recent and deeper past, which we comment on further in the discussion.

Previous literature has established a link between ROH patterns in a population and demography (Ceballos et al. 2018), specifically reflecting differences between large numbers of shorter ROH and fewer regions with long ROH. To explore this, we compared the normalized *N*_ROH_ to *F*_ROH_ for all individuals which had an *F*_ROH_ of at least 1%, as in (Ceballos et al. 2018). There were high levels of overlap in the space occupied by all species regardless of IUCN status (Fig. 2). The lines of best fit for Least Concern and Endangered species have steeper slopes compared to other species, indicating that at low *F*_ROH_ values they might only have a handful of long ROH, but that at high *F*_ROH_ values they are more likely to have many short ROH, but these differences are not significant with an ANOVA test (p-value = 0.68). When including captive-sampled individuals, a similar result is found, with the only notable results being Critically Endangered species overall having a high *N*_ROH_ across all *F*_ROH_ values (Fig. S4).

**Fig. 2.**
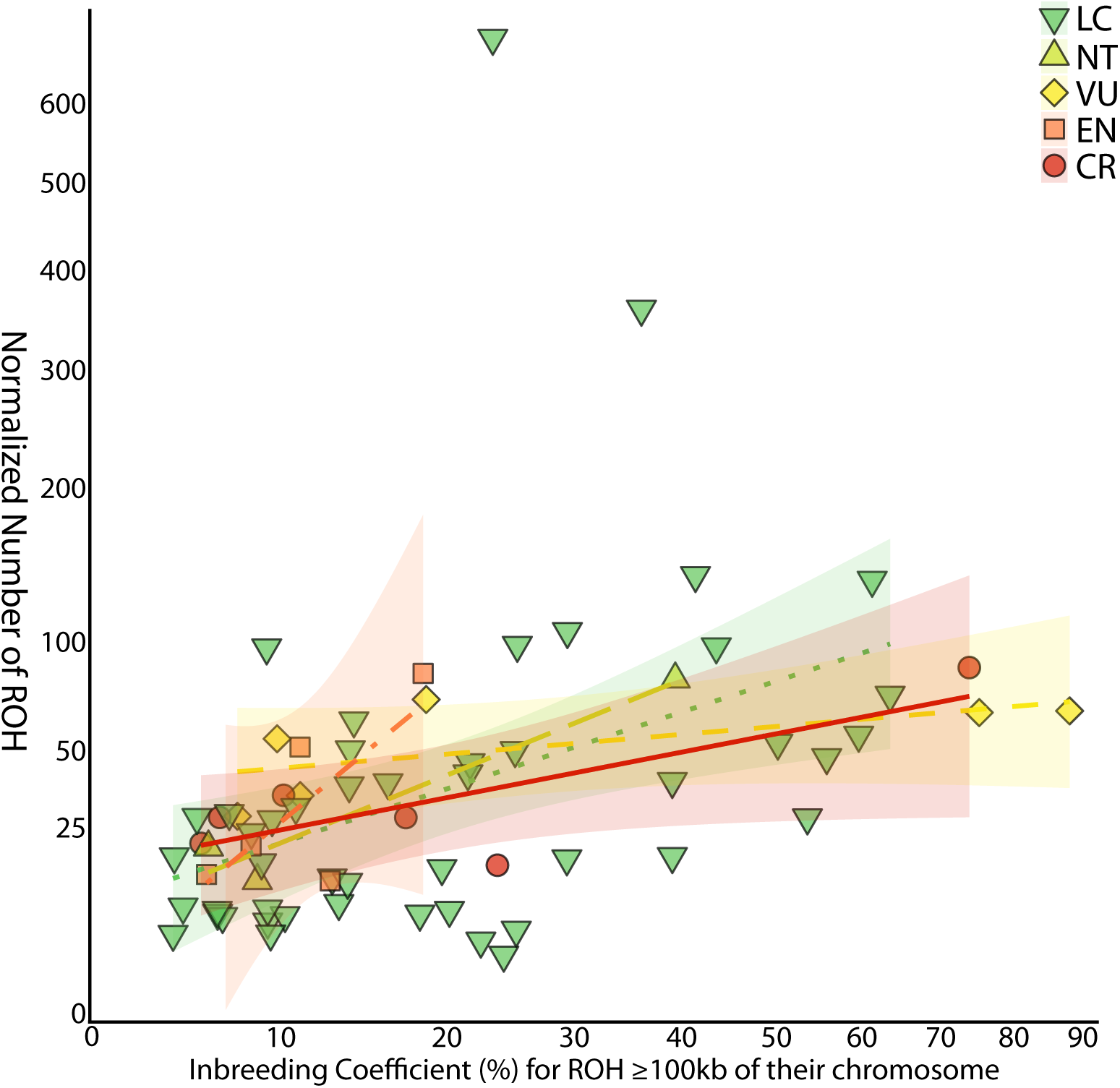
All wild individuals with an *F*_ROH_ of at least 1%, with their inbreeding coefficient and total count number of ROH compared (normalized by the number of chromosomes), and individuals color coded by IUCN status. LC, NT, VU, EN, CR denote IUCN status as in Figure 1.

### 3.2 Effective population size estimation through time with MSMC

We ran MSMC2, a sequentially Markovian coalescent method (Schiffels and Wang 2020), to estimate the effective population size (*N_e_*) through time for all species. Due to the lack of widespread life-history data for non-model and poorly studied vertebrates, generation time was not found for the majority of species and time is instead reported in number of generations. Given the wide range of generation times amongst known vertebrates (e.g. from 3 months for the house mouse to 24+ years in African elephants (Phifer-Rixey and Nachman 2015; Wittemyer et al. 2013)), establishing this information and studying changes in *N_e_* in years could allow for new understandings of vertebrate population dynamics and potential identification of major ecological events which impacted a wide variety of species.

We find that all categories decline in *N_e_*over recent time, and that beyond 100,000 generations they all have overlapping *N_e_* trajectories (Fig. 3). The consistent decline in the median *N_e_* value across groups is likely a result of MSMC assuming panmixia when in fact there is population structuring affecting *N_e_* values. However, in the most recent tens of thousands generations, we see some divergence by IUCN status: Least Concern species have a relatively steady *N_e_* that is higher than the other categories, while Critically Endangered and Endangered species have the lowest recent *N_e_* values (Fig. 3A).

**Fig. 3.**
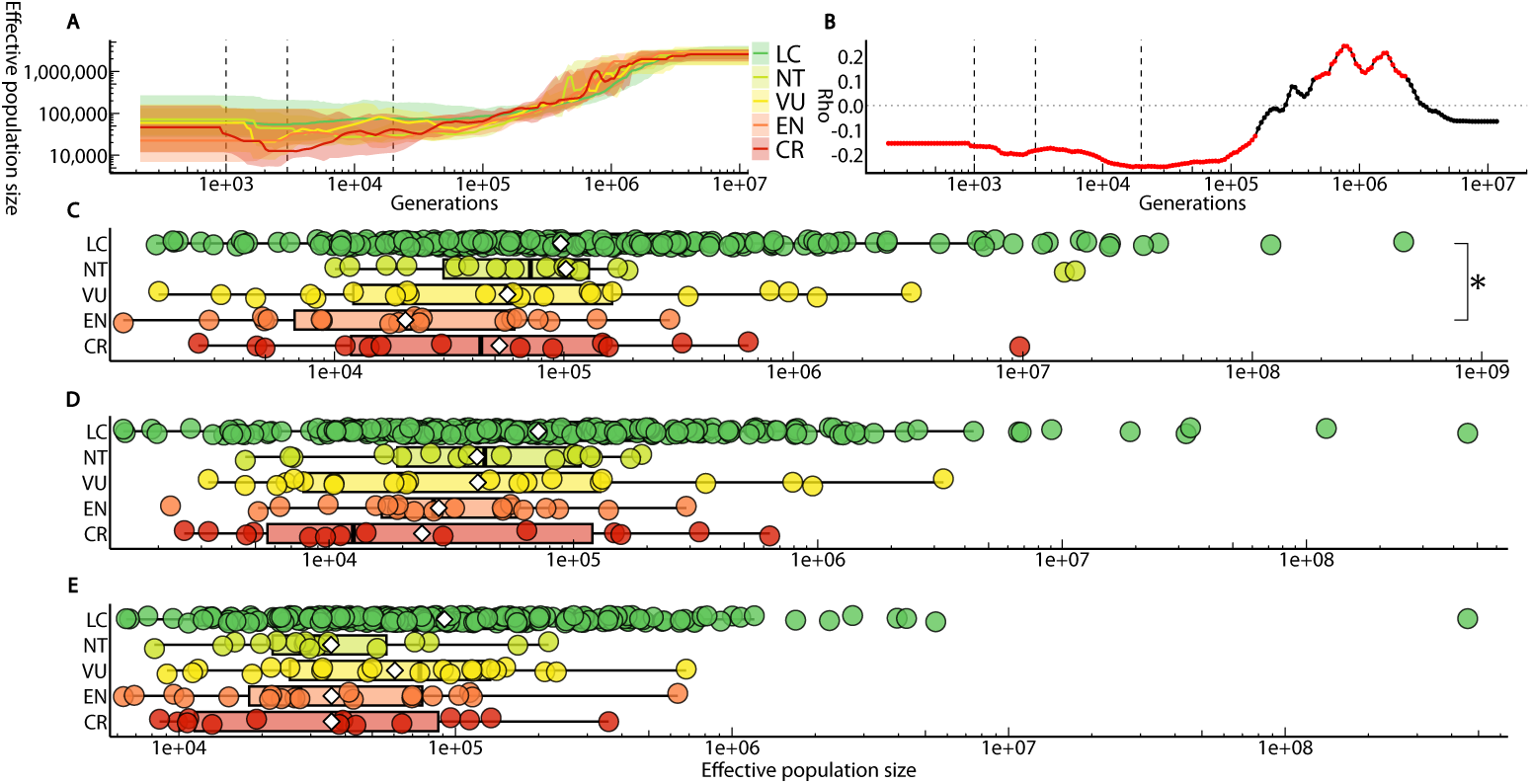
A) Effective population size over the last several million generations for wild species, grouped by IUCN status, with the line indicating the median and the shading indicating the 5 to 95 percentile region. Vertical dashed lines indicate the sampling time for the most recent time bin, 3,000 generations ago, and 20,000 generations ago. B) The Spearman rank correlation coefficient calculated at each time bin. Red indicates where the p-value is statistically significant, below 0.05. C) Effective population size estimated in the most recent time bin back to approximately 1,000 generations ago for wild species, grouped by IUCN status. D) Effective population size estimated approximately 3,000 generations ago for wild species, grouped by IUCN status. E) Effective population size estimated approximately 20,000 generations ago for wild species, grouped by IUCN status. LC, NT, VU, EN, CR denote IUCN status as in Figure 1.

We calculated the correlation between *N_e_* and IUCN status across all time bins using the Spearman rank correlation coefficient (Fig. 3B). We then compared the most recent time bin, which goes back to approximately 1,000 generations ago to two past time points, one corresponding to the time bin that contains 3,000 generations ago, and another at the time bin containing 20,000 generations ago. When looking at the recent past (1,000 generations ago), we see that all threatened categories have mean and median values lower than those for Least Concern and Near Threatened species, with Endangered species having the lowest recorded values (Fig. 3B). No species has a modern *N_e_* estimate less than 1,000, with the lowest observed contemporary *N_e_* in the Tasmanian devil (*Sarcophilus harrisii*) at 1,171 individuals. When including captive individuals, the results are broadly similar with the exception of Critically Endangered species now having the lowest median *N_e_*(Fig. S5). Compared to contemporary *N_e_*, there is a stronger relationship between *N_e_* and IUCN status 3,000 generations ago, both when looking at only wild species and when including captive individuals (Figs. 3C, S**??**). This trend strengthens further 20,000 generations ago. Through Bayesian regression while accounting for phylogenetic relationships, we see a significant relationship between *N_e_* and IUCN status across all three time points, but the effect size and confidence in the relationship increases as we go further back in time (Table 1).

### 3.3 Additional factors influencing genetic diversity measures

Given that population structure and gene flow can change with environment, we looked at whether ROH content varies by the type of IUCN habitat a species occupies. We found that species in marine aquatic habitats have significantly lower *F*_ROH_, both for all ROH lengths and also just for long ROH (Figs. S6-S7). Species that occurred in at least one marine habitat were flagged and compared to non-marine species for *F*_ROH_ content (Fig. 4A), and found to have significantly lower levels of ROH even when accounting for phylogenetic relationships using Phylolm and MCMCGlmm (p-value = 0.002 and p-value = 0.028 respectively). This same trend is found when looking at all ROH instead of long ROH, and when including captive individuals (Figs. S8-10). Both marine and non-marine species have large right skew, with positive outliers that have very high levels of *F*_ROH_, indicating a wide range of possible *F*_ROH_ values regardless of environment.

**Fig. 4.**
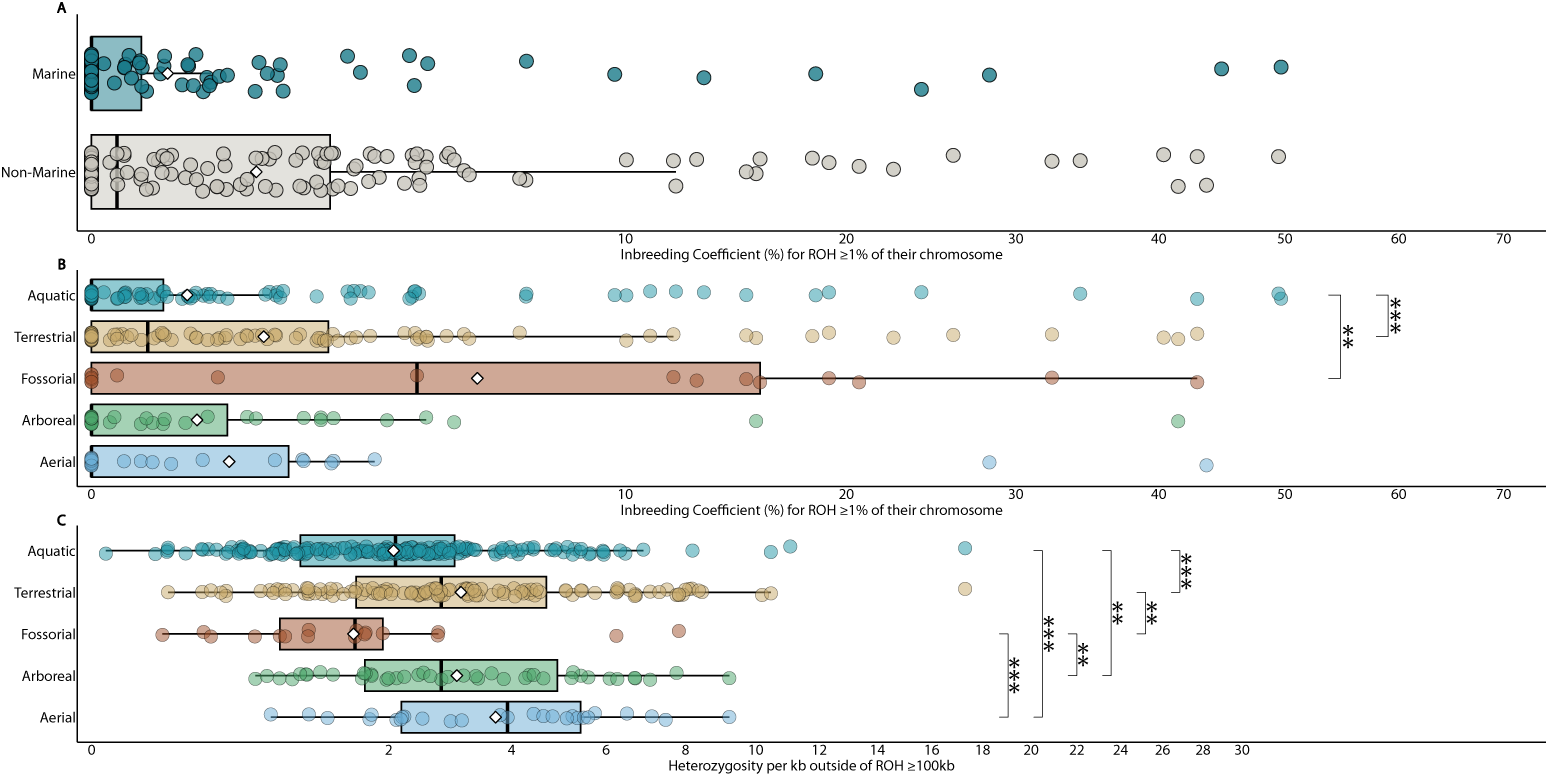
Genetic diversity and homozygosity as it relates to habitat for wild sampled species. A) The inbreeding coefficient for all ROH *≥* 1% of their chromosome in length for species that occur in at least one of the major marine habitats according to the IUCN. B) The inbreeding coefficient for all ROH *≥* 1% of their chromosome in length, grouped by strata (TetrapodTraits ‘microhabitat’). C) Heterozygosity in non-ROH regions (for ROH *≥* 100kb), with species grouped by strata. Aqu, aquatic; Ter, terrestrial; Fos, fossorial; Arb, arboreal; Aer, aerial.

We next investigated the more specific aspect of which stratum of the habitat is typically used by the species, provided by the TetrapodTraits microhabitat field as one or more per species of: aerial, aquatic, arborial, fossorial (digging and burrowing), and terrestrial (Moura et al. 2024). We found that aquatic species tend to have the lowest *F*_ROH_ values, followed by arboreal and aerial species, while fossorial species have the highest observed *F*_ROH_ values (Fig. 4B). This trend held when including captive-sampled species, and when looking at all ROH (Figs. S11-13). A Kruskal-Wallis test found significantly different values between strata for long ROH (p-value = 9.5e-05), and when using a Dunn’s test for pairwise comparisons, we find a significant difference between aquatic species and both terrestrial and fossorial species (Fig. 4B).

When we compare heterozygosity by stratum, we find the highest median and mean heterozygosity values in aerial species, and the lowest observed in fossorial species (Fig. 4C). This result is also consistent when including captive-sampled species (Fig. S14). A Kruskal-Wallis test shows significant differences in values between groups (p-value = 7.2e-07), and a Dunn’s test for pairwise comparisons shows a mix of significant differences between groups, including aquatic species having significantly lower heterozygosity than aerial, arboreal, and terrestrial species (Fig. 4C). These results suggest that life-history can impact genetic diversity and gene flow.

We next used mixed effects generalised least square (GLS) linear regression models to test the relative strength of the correlations between these different variables reported above and genetic diversity while accounting for phylogenetic relationships, with one model to predict *F*_ROH_ for long ROH and another for heterozygosity outside of ROH (Fig. 5). Heterozygosity is better explained than ROH, with 53.1% of its variance explained (adjusted *R*^2^ of the GLS model), while only 36.3% of *F*_ROH_ variance is explained in its model. Potentially this can be explained by stochasticity in mating patterns having a greater influence on ROH than on heterozygosity outside regions of ROH, which more reflects long term demography. Furthermore, about one third of the heterozygosity predictability comes from the habitat variables (19.7% adjusted *R*^2^ when including only habitat variables) whereas two thirds of the *F*_ROH_ predictability comes from habitat variables (24.1% adjusted *R*^2^).

**Fig. 5.**
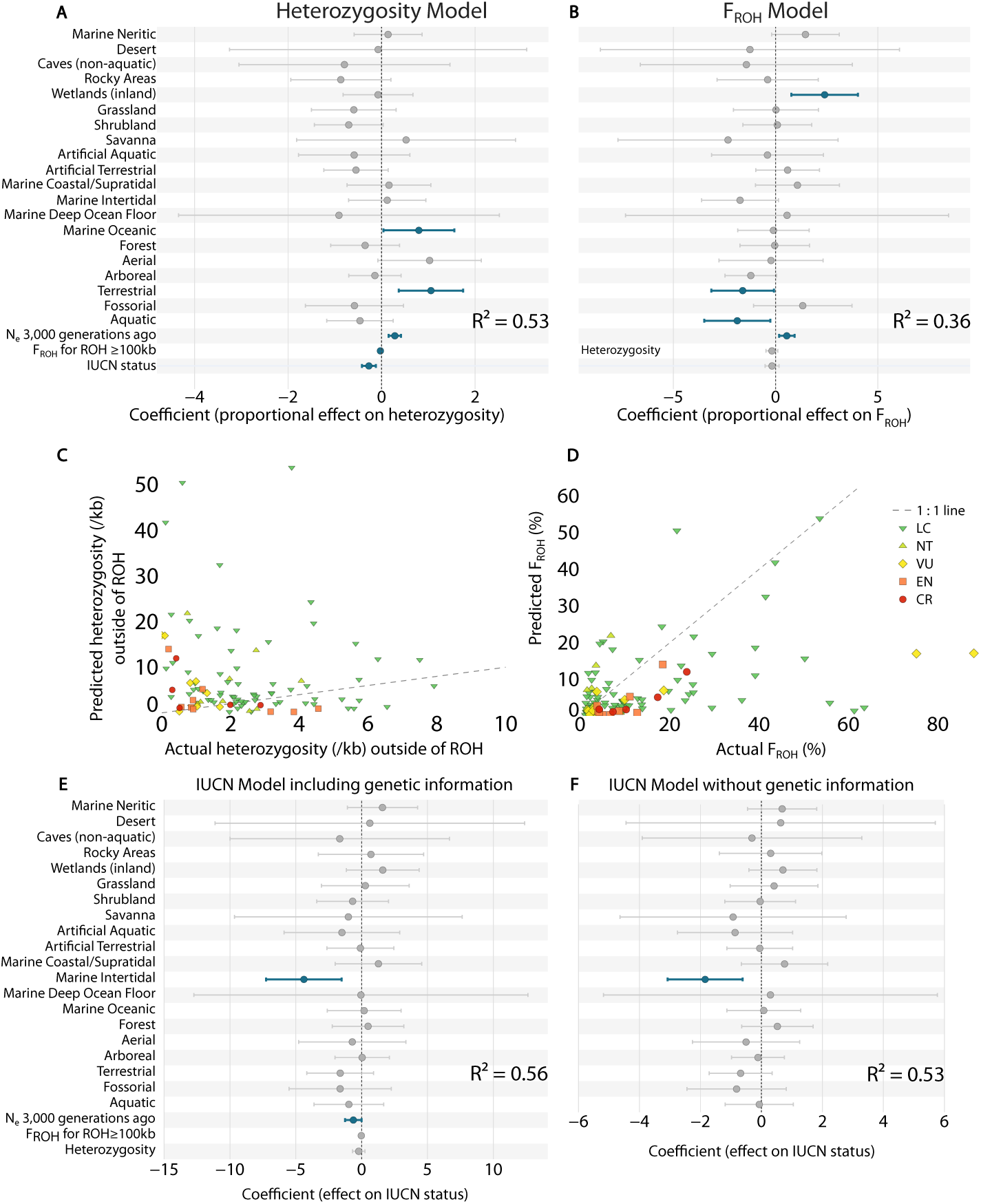
Mixed effects linear models to predict genetic diversity and IUCN status in wild-sampled species. (A) Coefficients and adjusted *R*^2^ for GLS model predicting heterozygosity per kb outside of ROH. (B) Same for model predicting *F*_ROH_ values for ROH *≥* 1% of their chromosome. (C,D) Actual versus predicted genetic diversity values for the models in A, B. The dashed line indicates a perfect 1:1 linear relationship. (E,F) Mixed effects linear models to predict IUCN status in wild-sampled species (E) Coefficients and adjusted *R*^2^ for GLS model predicting IUCN status, with genetic information included (F) As in (A) with only habitat and strata included. Bars in (A,B,E,F) indicate 95% confidence intervals. Significant coefficients (p *<* 0.05) are highlighted in blue. LC, NT, VU, EN, CR denote IUCN status as in Figure 1.

When looking at the proportional contribution of different factors, we see that being terrestrial or living in the marine oceanic habitat are positively linked with heterozygosity outside of ROH, while living in shrubland or having higher IUCN status are negatively linked (Fig. 5A). Inbreeding is positively correlated with species found in wetlands, while terrestrial and aquatic species tend to have lower *F*_ROH_ (Fig. 5B). When looking at the predicted values for genetic diversity compared to those observed, we find that the models are primarily driven by Least Concern species, which are more numerous, and are poorer predictors for species in other categories (Fig. 5C-D).

Finally, we used a phylogenetically corrected mixed effects GLS model to predict IUCN status (as an ordinal scale from 0-4) of species based on their habitat and strata, *F*_ROH_, heterozygosity outside of ROH regions, and historical effective population size 3,000 generations ago (Fig. 5E). Together these explain 56% of the variance in IUCN status. We see that the two factors with independently significant contributions to this model are that species in Marine Intertidal habitats are likely to be in a lower IUCN category, and a small but significant effect from historical *N_e_* 3,000 generations ago, with higher historical effective population size corresponding to lower current IUCN status. However, overall most of the predictive power comes from the non-genetic factors: removing genetic information removes only approximately 3% of the adjusted *R*^2^ value (53% versus 56%), demonstrating that habitat variables provide the majority of the predictive power in this model (Fig. 5F).

IDRisk is a new metric introduced to predict risk of inbreeding depression if the population should decline, based on the product of *F*_ROH_ and heterozygosity outside of ROH (Kyriazis et al. 2025a). The underlying idea is that species with both high heterozygosity and high *F*_ROH_ are more vulnerable to homozygozing recessive deleterious variants. When looking at IDRisk in species with *F*_ROH_ for long ROH of at least 5%, we found almost all species with predicted high or extreme high IDRisk are listed as Least Concern, whereas threatened species are either at low or moderate risk (Fig. 6). This appears to be due to threatened species with high inbreeding levels having low heterozygosity in non-ROH regions or vice versa. This result perhaps reflects that IDRisk is not intended to be a direct measure of current extinction risk, but rather a conditional measure for if circumstances were to change. When including captive-sampled species, we find the trend remains largely the same, but with a few threatened species at high risk, and one at extreme risk (Fig. S15).

**Fig. 6.**
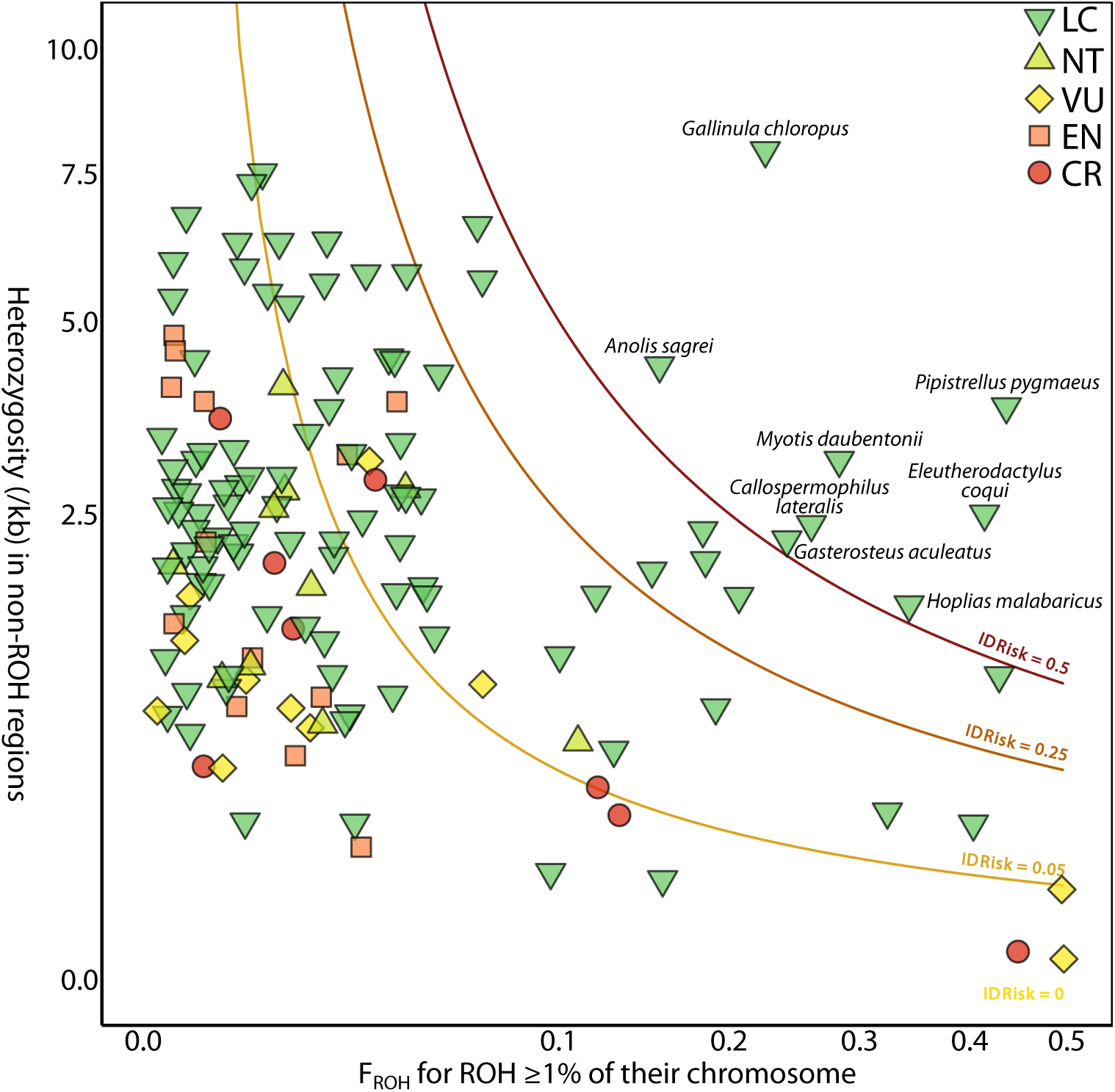
Inference of IDRisk (predicted risk of inbreeding depression if the population size were to decline, Kyriazis et al., 2025) for all wild-sampled species, with shapes categorizing IUCN status. Clines show the cutoffs for low to extreme IDRisk. Low risk is less than 0.05, moderate risk is between 0.05 and 0.25, high risk is between 0.25 and 0.5, and extreme risk is above 0.5. LC, NT, VU, EN, CR denote IUCN status as in Figure 1.

## 4 Discussion

Given the current biodiversity crisis, it is important to understand factors which contribute to extinction risk in species and how we can effectively help species recover. This requires multidisciplinary work, including in understanding genetic factors. There is currently a lack of consensus about the importance of measures of genetic variation. Many state that low genetic variation across a population is an indicator of doing poorly while high overall variation is associated with good fitness and lower extinction risk (Reed and Frankham 2003; Theissinger et al. 2023; Willi et al. 2022; Kardos et al. 2021; Willoughby et al. 2015; Leigh et al. 2019; Romiguier et al. 2014), but others argue that the importance of neutral genetic diversity is exaggerated, citing that only a weak correlation exists between the genome-wide nucleotide diversity in a species and IUCN Red List status (Teixeira and Huber 2021; Doyle et al. 2015; Brüniche-Olsen et al. 2018, 2019; Grossen et al. 2020). Our results provide some support for both positions. We find that heterozygosity outside of runs of homozygosity significantly negatively correlates with IUCN status. However there is a high overlap in the ranges between categories, reflecting that the correlation is far from total (Fig. 1). Nevertheless, this supports the view that species with fewer variants in their genome are more likely to be threatened, suggesting that smaller long-term historical effective population size is a risk factor for extinction vulnerability.

It is striking that in our MSMC2 analysis the historical *N_e_* estimates a few thou-sand generations ago or tens of thousands of generations ago are better correlated with IUCN status than present-day estimated *N_e_*, with the estimate from 20,000 generations ago better correlated than heterozygosity. This suggests that longer term demography provides an important indicator of vulnerability, at least when looking at a single individual instead of population data. This could be because present-day estimates from SMC models are less accurate, but alternatively it may reflect that species that have suffered from low *N_e_* for a substantial period of time are more vulnerable to sudden changes, for example due to anthropogenic habitat threats. These match previous results found in mammals during the Zoonomia project (Wilder et al. 2023). Previous research proposed that a species needs an *N_e_* of at least 50 to avoid short term inbreeding depression, and at least 500 for retaining long-term evolutionary potential, known as the 50/500 rule (Souĺe and Wilcox 1980), with others arguing for a similar 100/1,000 rule (Frankham et al. 2014). While none of the species we analysed have *N_e_* estimations below 1,000 in the most recent time bin, we had eight species with *N_e_* estimations below 2,000, three of which were close to the 1,000 individual threshold. Given that the most recent time bin can span anywhere from 10s to 1000s of generations, it is possible that the contemporary *N_e_* estimates for these species are actually lower than is being predicted by MSMC2; indeed this is definitely the case for Rice’s whale (Aguilar-Gómez et al. 2026). As such, it is possible these species are below the thresholds for safely being able to retain their evolutionary potential in the long-term, and should be studied in closer detail to ensure proper management. We note that adding only a small number of independent sequenced samples can substantially improve recent *N_e_* estimates Schiffels and Wang (2020).

Traditionally, ROH have been extensively studied in humans and economically important species such as livestock (see (Sams and Boyko 2019; Nothnagel et al. 2010; Selli et al. 2021; Tao et al. 2025; Pegolo et al. 2025; Mota et al. 2024; Fatma et al. 2023)), but they have recently begun to be explored in wild populations in the context of conservation and management (see Stanhope et al. (2023); Hewett et al. (2023b); Stoffel et al. (2021); Foote et al. (2021); Kardos et al. (2018)). Most studies quantified ROH length in bases, with long ROH typically defined as 1-2Mb or longer. This is fine within species, but between species we expect ROH lengths to vary inversely with recombination rate, and hence approximately proportionally with chromosome size. The vertebrate genomes sequenced by the VGP range in size by over two orders of magnitude, from 324Mb for the Greater Pipefish (*Syngnathus acus*) to 40.5Gb for the West African Lungfish (*Protopterus annectens*), with a similar range of chromosome sizes, making use of absolute lengths inappropriate. We addressed this by normalizing ROH size by the length of the chromosome it is found on, with every ROH ranging from 0-100%, allowing for comparative ROH studies across a wide range of vertebrate species. We suggest standardizing on this in guidelines for analysis of inbreeding across species.

Building on this, we found that our samples from more threatened species are more likely to have related parents, as indicated by the positive correlation between *F*_ROH_ and IUCN status. Similar to heterozygosity, there was also in this case a high overlap in ranges between categories, indicating a wide range of inbreeding patterns are possible regardless of how threatened a species is. We discuss as particular examples the European fallow deer (*D. dama*), the European badger (*M. meles*), and Rice’s whale (*B. ricei*).

The fallow deer had a bimodal distribution of ROH with both short and ultralong ROH (Fig. 1C). This is indicative of smaller populations or a population bottleneck, and also that the parents of the individual were closely related. Fallow deer are found throughout the majority of Europe, having been spread by humans from their native range in Turkey over the past two millennia (Masseti and Mertzanidou 2008). The individual sampled was from the wild in Denmark, however populations throughout Europe are almost never free-ranging and instead managed in forested parks, and as such the habitat fragmentation and lack of gene flow are consistent with both high levels of recent inbreeding and small *N_e_* over the last tens to hundreds of generations (Masseti and Mertzanidou 2008).

A similar story is found in the European badger. The individual sampled had a lower inbreeding coefficient (39.1%), but the majority of ROH are ultralong (Fig. 1C). This is indicative of inbreeding between very closely related parents. European badgers live in large social groups centred around their dens, which increases the likelihood of consanguinity, particularly in fragmented habitats (Dugdale et al. 2008; Annavi et al. 2014). In this specific case, the badger sampled, individual 1581, came from a population in Wytham Woods, Oxfordshire, UK, which have been studied in detail for over 30 years (Newman et al. 2022). He was part of a group whose pedigree is being tracked, and has been confirmed to be the result of an uncle-niece pairing (P. Holland, personal communication, November, 2025).

By comparison, Rice’s whale (*B. ricei*) is a species with a very high *F*_ROH_ at 73.8%, but has an approximately normal distribution of ROH sizes from small to large (Fig. 1C). This indicates that the individual sampled was likely from a small population, leading to moderate levels of parental relatedness. This matches the known distribution of this species. Rice’s whale is only found in the Gulf of Mexico, and as of 2018 is estimated to have only approximately 50 individuals left (Rosel et al. 2022). Indeed, recent research has shown that low heterozygosity and high levels of ROH are present across Rice’s whales, and that they have existed as a notably small population for thousands of years (Aguilar-Gómez et al. 2026). Here there is a concern that their reduced genetic diversity might restrict their ability to adapt to changing environmental conditions.

Given the overlap in genetic diversity ranges across IUCN categories, we examined other factors which could influence diversity such as habitat and stratum, which is the part of a habitat that a species primarily occupies or forages in. Fossorial species (those that burrow and dig) had the highest levels of inbreeding, and the lowest heterozygosity (Fig. 4B-C). As we saw in the European badger (*M. meles*), fossorial life history traits can lead to frequent consanguinity (Dugdale et al. 2008; Annavi et al. 2014). Similar to the badger, other fossorial species typically have lower dispersal and smaller, patchy ranges, leading to higher levels of population structure and less gene flow, which can contribute to lower heterozygosity and higher levels of inbreeding (Reuber et al. 2024). At the other extreme, aerial and arboreal species had the highest reported heterozygosity on average, together with amongst the lowest *F*_ROH_ values behind aquatic species (Fig. 4B-C). For birds, this is likely due to many species having higher dispersal and less population structure (Kozakiewicz et al. 2018).

Species in marine habitats, and more broadly aquatic species, show a more mixed picture. They tend to show lower levels of inbreeding in comparison to other species (Fig. 4A-B), but also lower heterozygosity across all Aquatic species (Fig. 4C). This contrasts with a previous study which found that aquatic chordates have higher genetic diversity than their terrestrial counterparts (Leffler et al. 2012). However, other studies have reported that teleosts, the largest group within ray-finned fishes, tend to have lower genetic diversity than other major vertebrate groups (Ward et al. 1992), and that marine fishes tend to have higher levels of genetic diversity in comparison to freshwater fishes due to higher levels of gene flow (Martinez et al. 2018). These patterns at least partly reflect variation in gene flow, with studies showing that marine aquatic species can have substantial amounts of gene flow despite retaining fine-scale population structure, in at least some cases due to larval dispersal even when adult movement is restricted due to environmental barriers such as currents or temperature gradients (Banks et al. 2007; Pascual et al. 2017; Cooke et al. 2016).

Overall, approximately one third to one half of genetic diversity in an individual’s genome can be explained by their life history and IUCN status (Fig. 5), showing the stochasticity and wide range of factors that go into heterozygosity, ROH formation, and subsequent shortening through recombination. Despite noise, the model is informative - we find that being terrestrial or aquatic, and specifically living in marine oceanic habitat, is correlated with increased genetic diversity, while being fossorial or living in shrubland is correlated with decreased genetic diversity (Fig. 5). When predicting IUCN status using combined ecological and genetic information, we find that ecological information provides most of the predictive power, with a small but significant impact from genetic information, interestingly best captured via the inferred historical effective population size rather than heterozygosity or *F_ROH_* (Fig. 5E,F). When looking across mammals, the Zoonomia consortium found similar results, with ecological data being more predictive of IUCN status than genetic (Wilder et al. 2023). In conclusion, our results across a wide range of vertebrates support the view that ecological context is a more important factor associated with extinction risk in our current, anthropogenic era than genetic species diversity. Much of the association seen between IUCN status and measures of genetic diversity, such as heterozygosity and the inbreeding coefficient derived from runs of homozygosity that we focus on in this study, can be explained by association of genetic diversity to the ecological habitat parameters. Nevertheless, we do see a small but significant independent contribution of genetic information to risk prediction. Interestingly, this is better captured by historical effective population size inferred from demographic models several thousand generations ago, rather than current diversity, consistent with the notion that longer term lower diversity puts populations more at risk when subject to anthropogenic shock.

## 5 Data Availability Statement

All sequence data analysed are available at NCBI under the accession numbers given in Supplementary Data. All scripts used in analysis for both the VGP data analysis, along with a list of all packages used in the analyses, are available at https://github.com/alwaysamanda/VGP Phase1 Heterozygosity FROH analysis/tree/main. Scripts were written using Python (https://www.python.org) and R (https://www.r-project.org).

## Supporting information

VGP Phase I Consortium Extended Author List

## Acknowledgements

We would like to acknowledge funding from the Wellcome Trust to R.D. (award 317408/Z/24/Z) and Christ’s College Cambridge for support to A.G. We thank Minoli Daigavane, Runyang Nicolas Lou, and Peter Sudmant for curating and sharing data on captivity status, along with Heng Li for discussions about early stages of this work and MSMC analysis. We thank Peter Holland for communication about the Wytham Woods Badger project and providing lineage data for the individual sampled. Finally, we express our deep gratitude to all the providers of samples to be sequenced and those who generated the genome assemblies that we have used. For the purpose of open access, the author has applied a CC BY public copyright licence to any Author Accepted Manuscript version arising from this submission.

## 6 Statements and Declarations

### 6.3 Author Contributions

AG and RD contributed to study conception and design. Sequence data was provided by the Vertebrate Genomes Project Phase I. Analysis and visualization was done by AG. The original draft was written by AG. Review and editing of the manuscript was done by AG and RD. RD supervised the project. Funding acquisition was done by RD.

