## Supplementary Figures for "The relationship of genetic diversity and inbreeding to extinction risk across over 500 vertebrate diploid genomes"

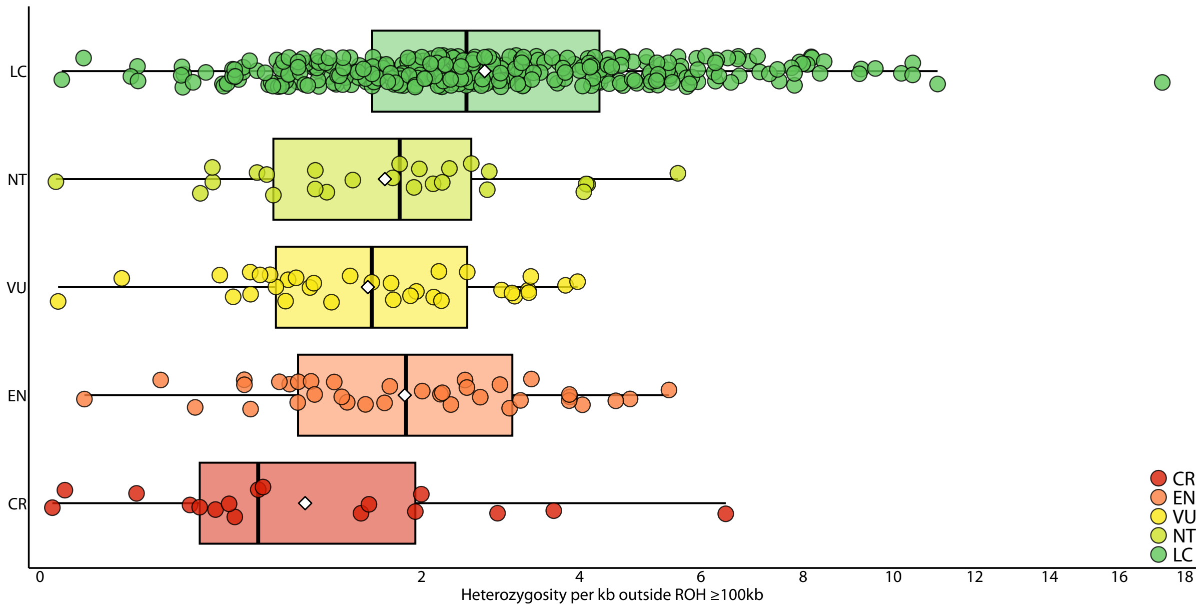


Figure S1: Heterozygosity outside of ROH (for ROH $\geq$ 100kb) for all species, including captive-sampled, grouped by IUCN status. LC, Least Concern; NT, Near Threatened; VU, Vulnerable; EN, Endangered; CR, Critically Endangered.

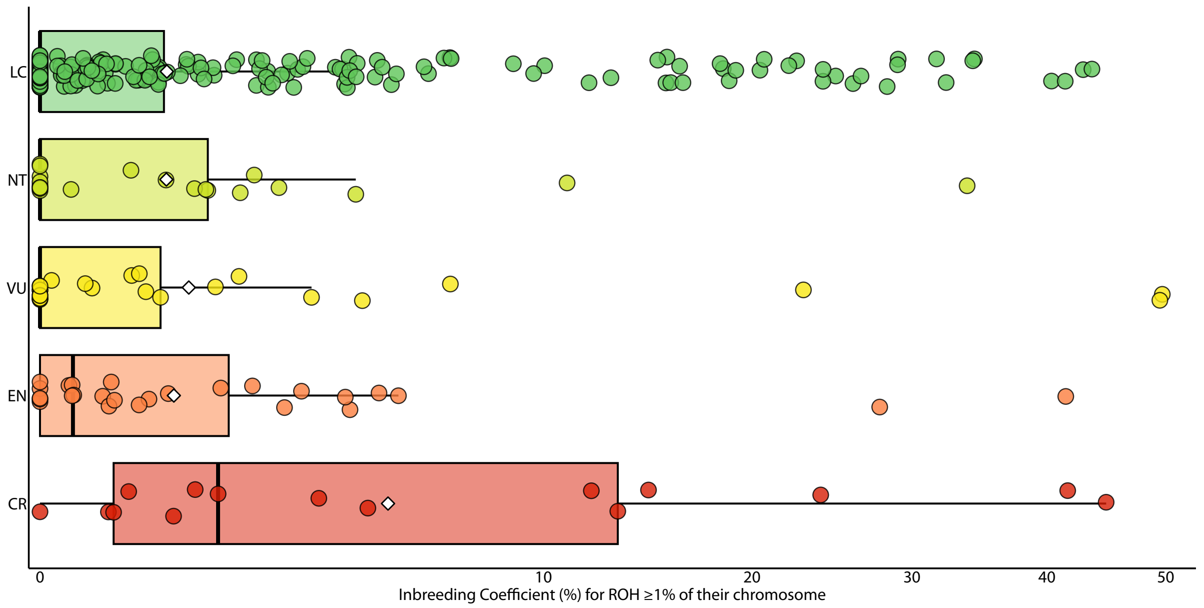


Figure S2: The inbreeding coefficient in all species, including captive-sampled for long ROH that are at least 1% of their chromosome in length. LC, Least Concern; NT, Near Threatened; VU, Vulnerable; EN, Endangered; CR, Critically Endangered.

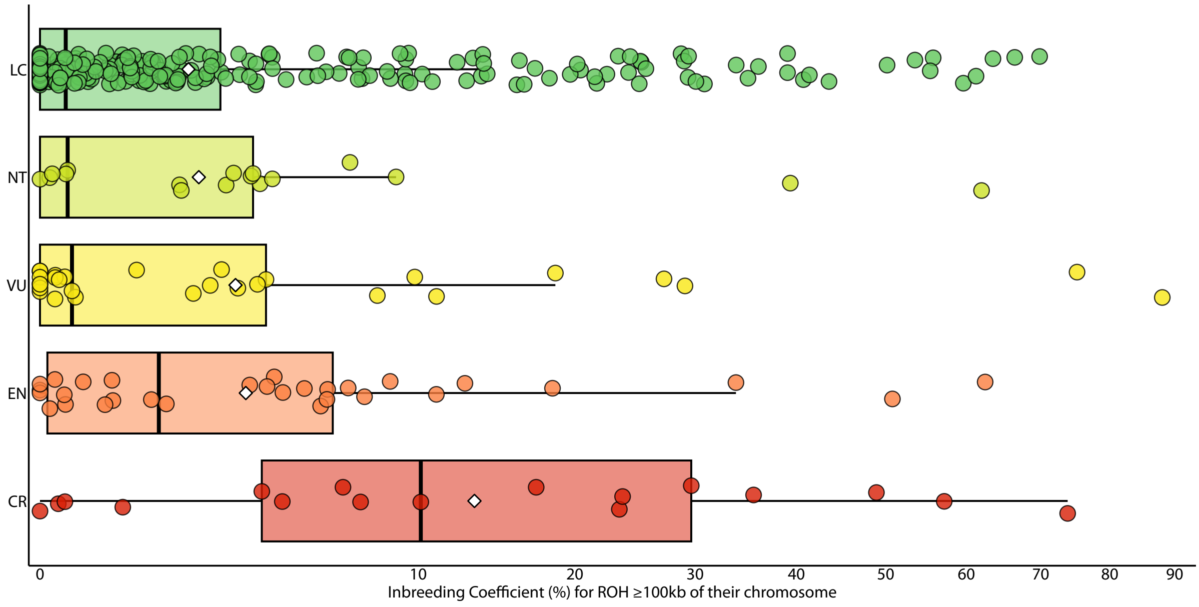


Figure S3: The inbreeding coefficient in all species, including captive-sampled for all ROH that are at least 100kb in length, grouped by IUCN status. LC, Least Concern; NT, Near Threatened; VU, Vulnerable; EN, Endangered; CR, Critically Endangered.

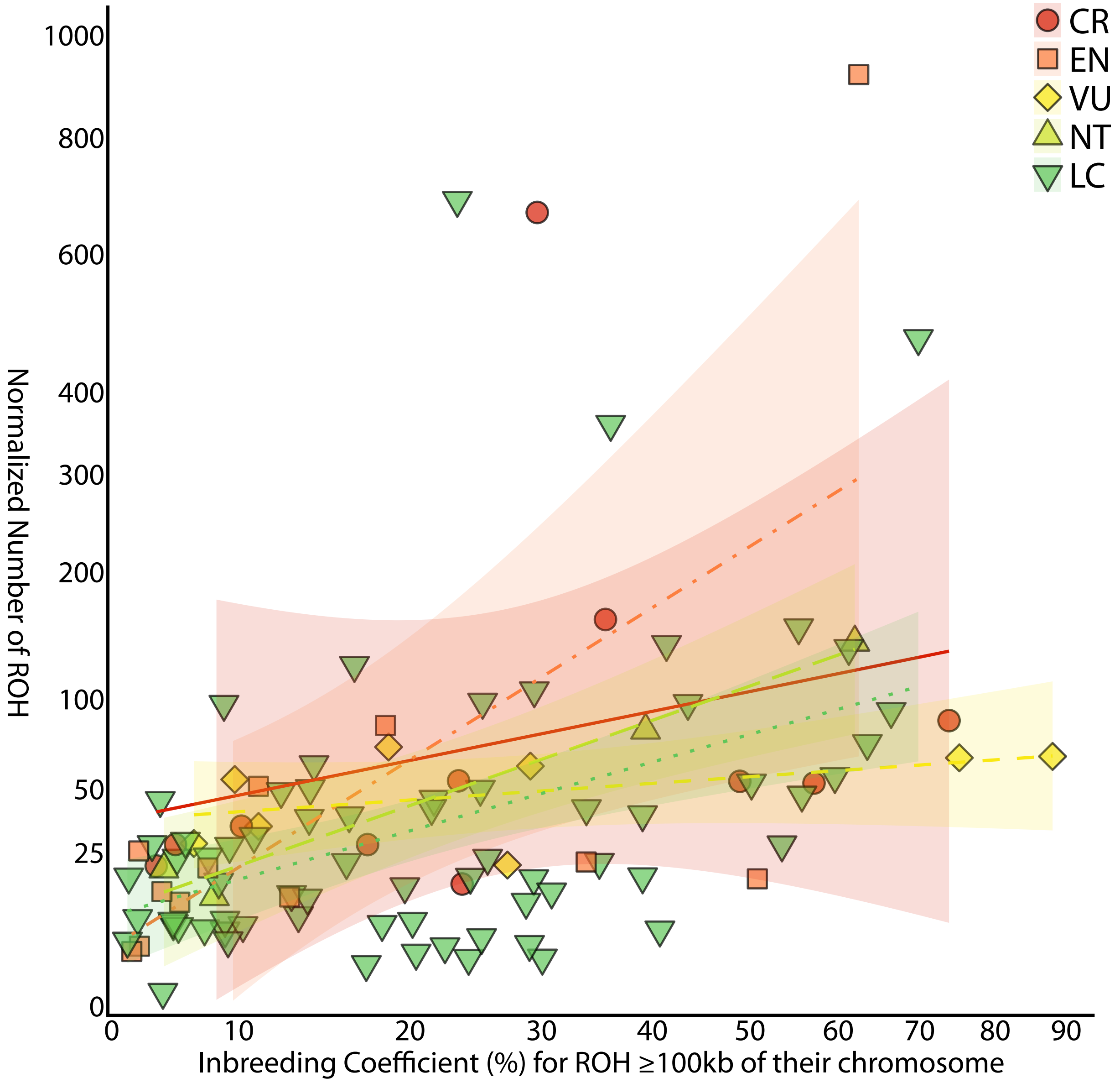


Figure S4: All individuals, including captive-sampled, with an $F_{\mathrm{ROH}}$ of at least 1%, with their inbreeding coefficient and total count number of ROH compared. Individuals are color-coded and have different shapes based on IUCN status. LC, Least Concern; NT, Near Threatened; VU, Vulnerable; EN, Endangered; CR, Critically Endangered.

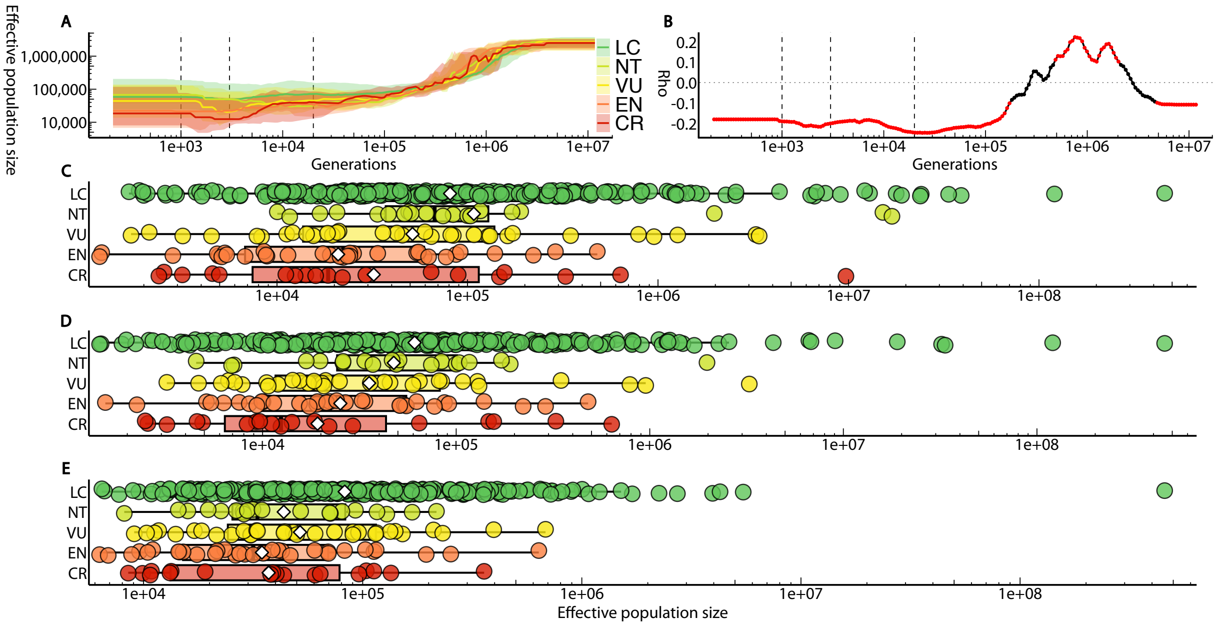


Figure S5: A) Effective population size over the last several million generations for all species including captive-sampled, grouped by IUCN status, with the solid lines indicating the median and the shading indicating the 5 to 95 percentile region. Vertical dashed lines indicate the sampling time for the most recent time bin, 3,000 generations ago, and 20,000 generations ago. B) The Spearman rank correlation coefficient calculated at each time bin. Red indicates where the p-value is statistically significant, below 0.05. C) Effective population size estimated in the most recent time bin back to approximately 1,000 generations ago for wild species, grouped by IUCN status. D) Effective population size estimated approximately 3,000 generations ago for wild species, grouped by IUCN status. E) Effective population size estimated approximately 20,000 generations ago for wild species, grouped by IUCN status. LC, Least Concern; NT, Near Threatened; VU, Vulnerable; EN, Endangered; CR, Critically Endangered.

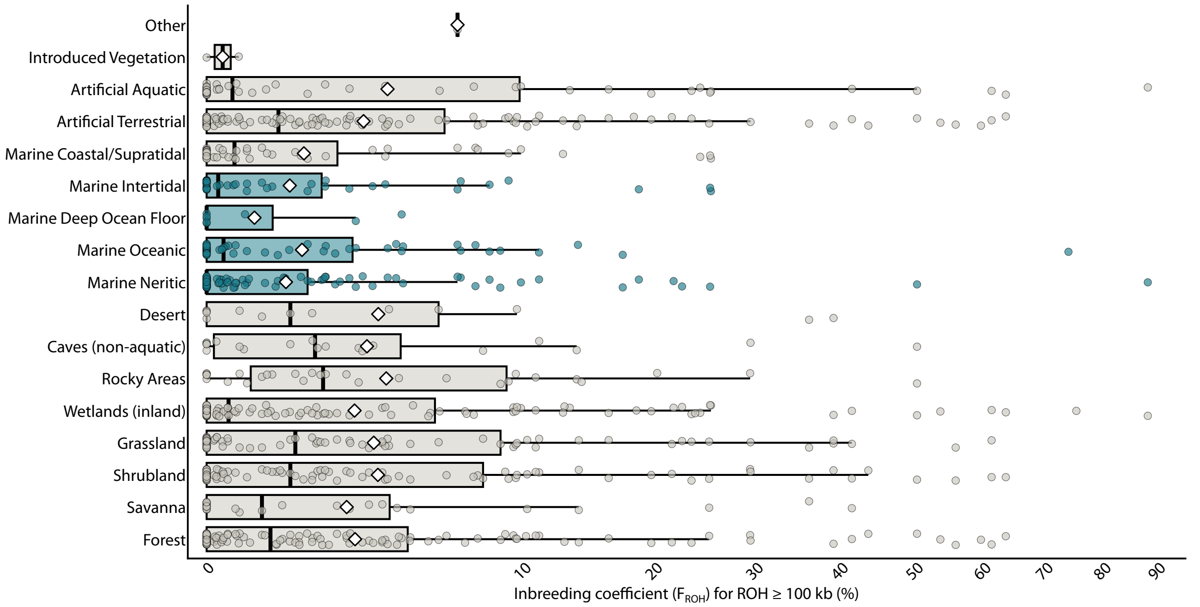


Figure S6: The inbreeding coefficient for all ROH $\geq$ 100kb for wild-sampled species, grouped by major IUCN habitat types. If a species occurs in multiple habitats, it is duplicated so that it appears once in each habitat. Marine habitats are highlighted in blue.

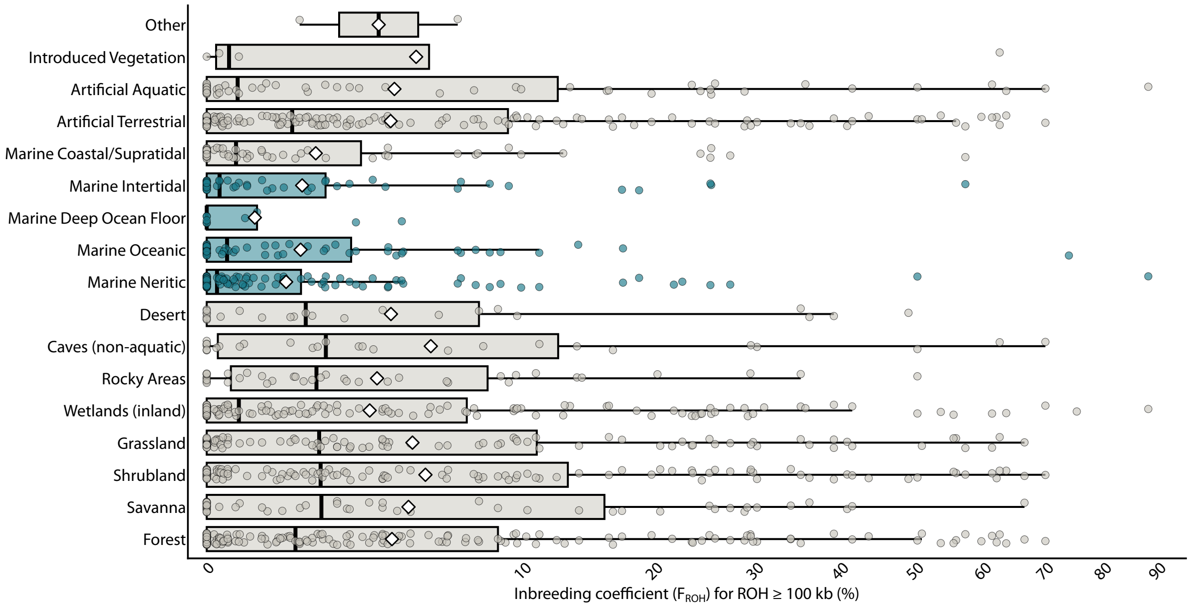


Figure S7: The inbreeding coefficient for all ROH $\geq$ 100kb for all species, including captive-sampled, grouped by major IUCN habitat types. If a species occurs in multiple habitats, it is duplicated so that it appears once in each habitat. Marine habitats are highlighted in blue.

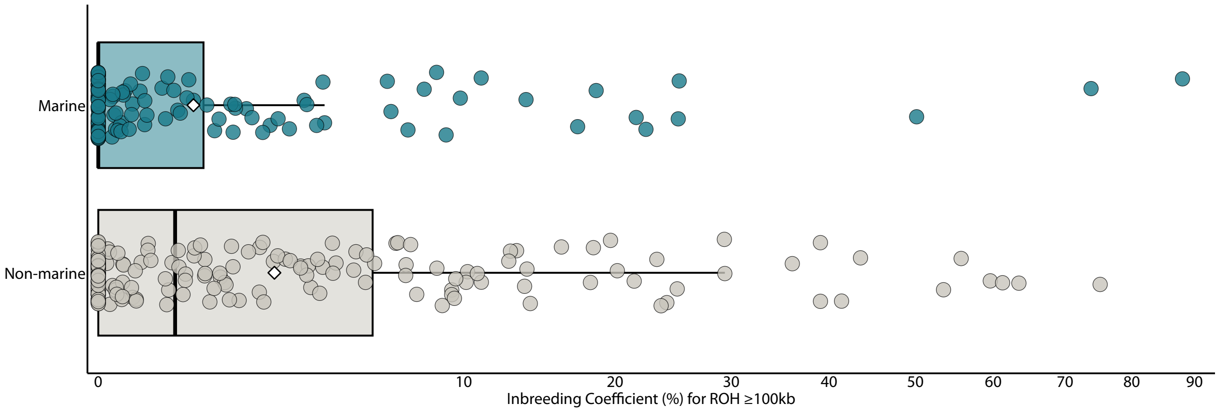


Figure S8: The inbreeding coefficient for all ROH $\geq$ 100kb for wild-sampled species, with species grouped by whether they occur in at least one of the marine aquatic habitats as defined by the IUCN major habiats.

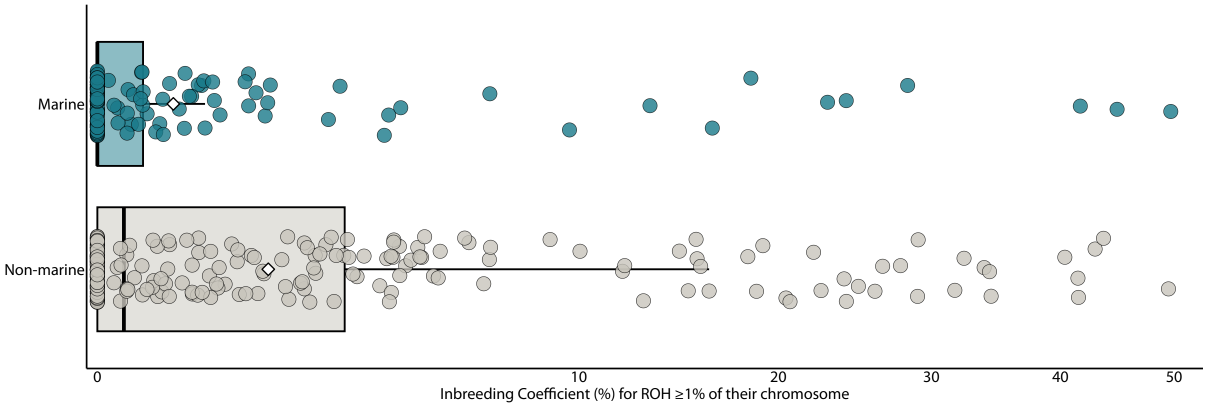


Figure S9: The inbreeding coefficient for all ROH $\geq$ 1% of their chromosome for all species, including captive-sampled, with species grouped by whether they occur in at least one of the marine aquatic habitats as defined by the IUCN major habiats.

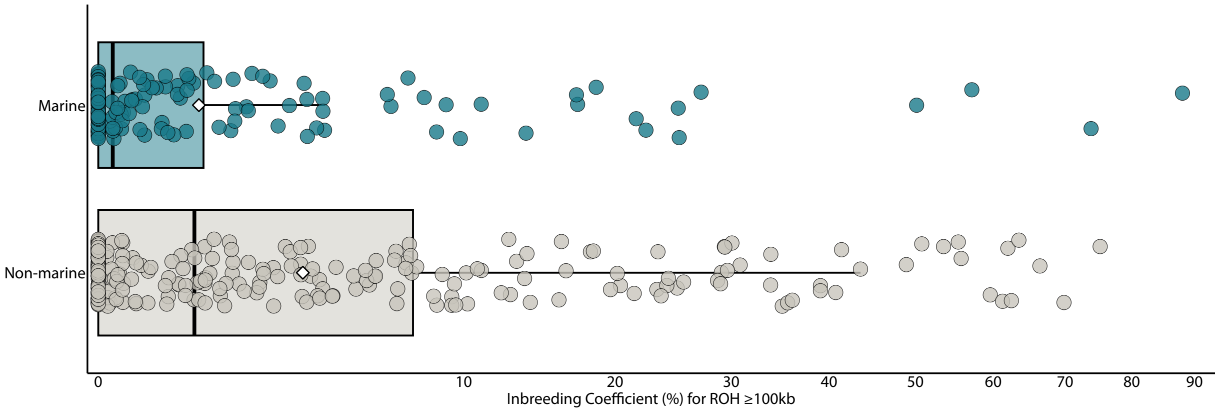


Figure S10: The inbreeding coefficient for all ROH $\geq$ 100kb for all species, including captive-sampled, with species grouped by whether they occur in at least one of the marine aquatic habitats as defined by the IUCN major habiats.

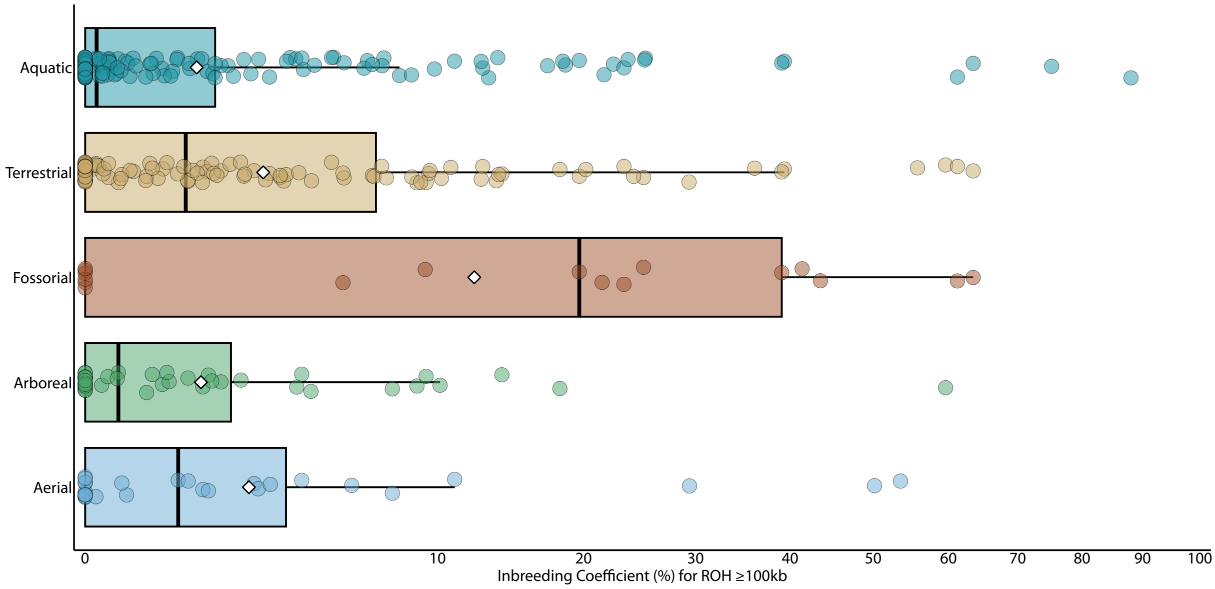


Figure S11: The inbreeding coefficient for all ROH $\geq$ 100kb for wild-sampled species, grouped by stratum. Aqu, aquatic; Ter, terrestrial; Fos, fossorial; Arb, arboreal; Aer, aerial.

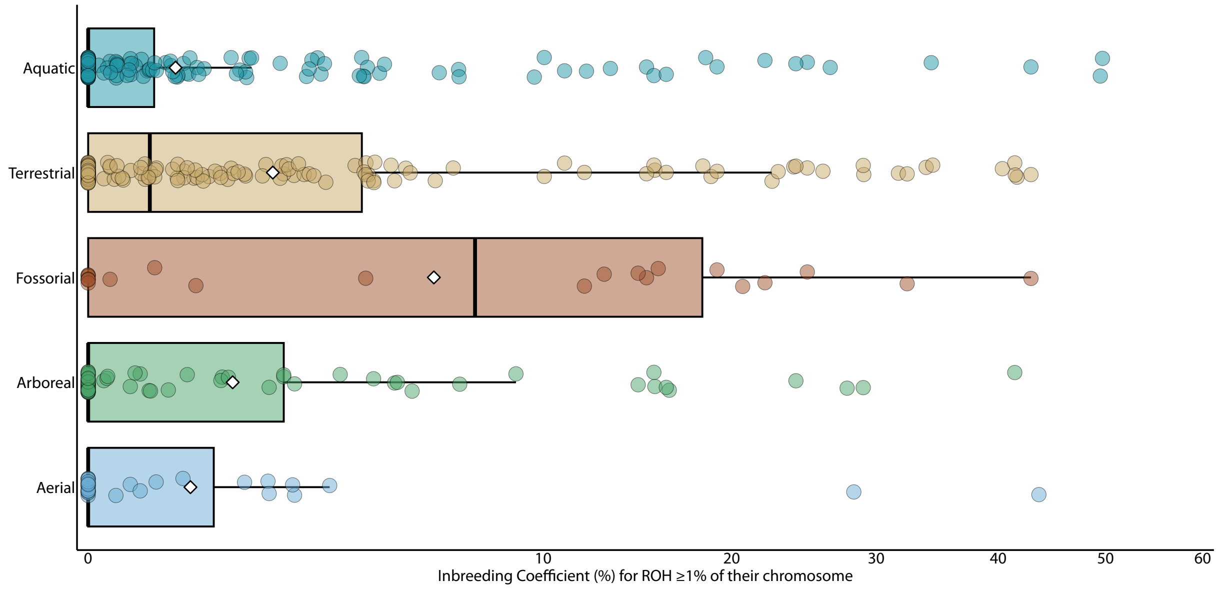


Figure S12: The inbreeding coefficient for all ROH $\geq$ 1% of their chromosome in length for all species, including captive-sampled, grouped by stratum. Aqu, aquatic; Ter, terrestrial; Fos, fossorial; Arb, arboreal; Aer, aerial.

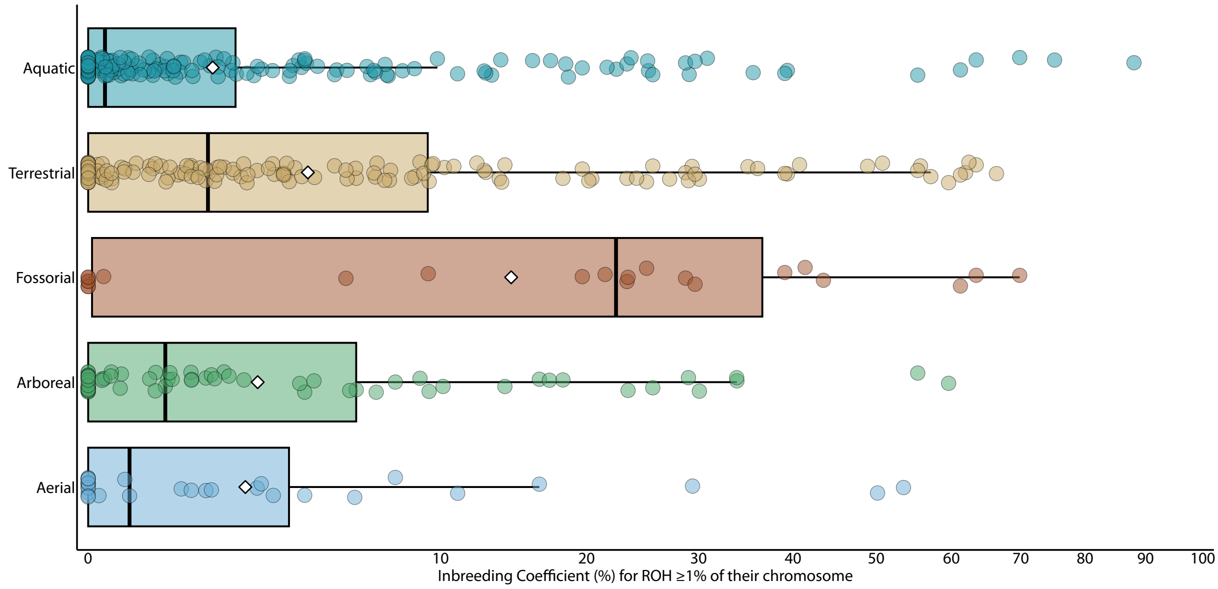


Figure S13: The inbreeding coefficient for all ROH $\geq$ 100kb for all species, including captive-sampled, grouped by stratum. Aqu, aquatic; Ter, terrestrial; Fos, fossorial; Arb, arboreal; Aer, aerial.

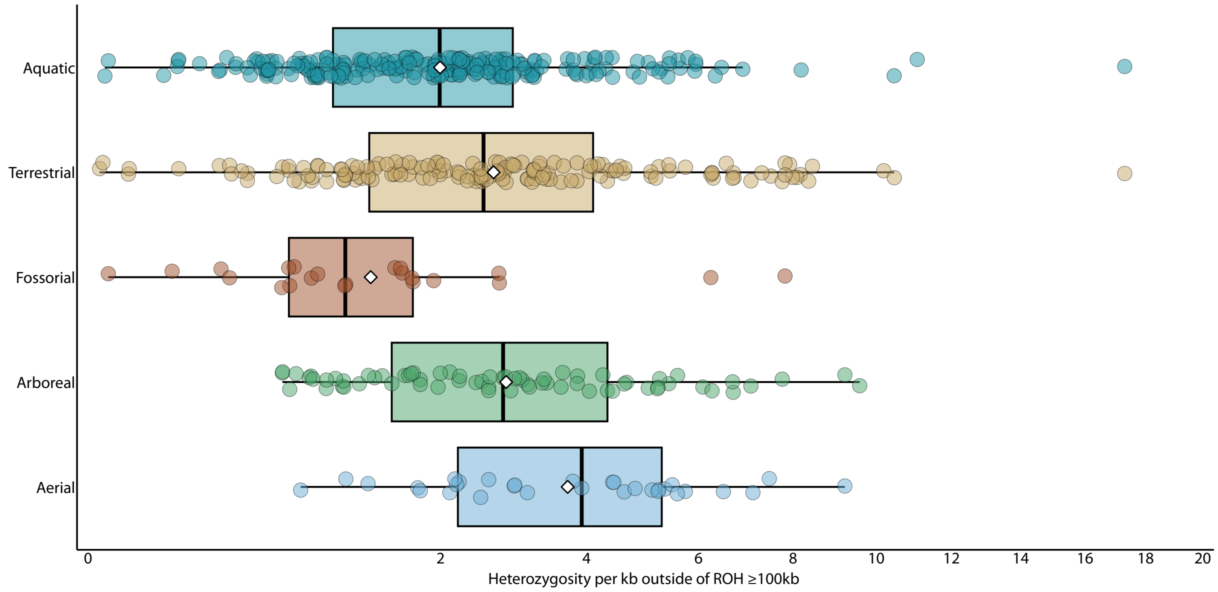


Figure S14: Heterozygosity per kb outside of ROH for all species, grouped by stratum. Aqu, aquatic; Ter, terrestrial; Fos, fossorial; Arb, arboreal; Aer, aerial.

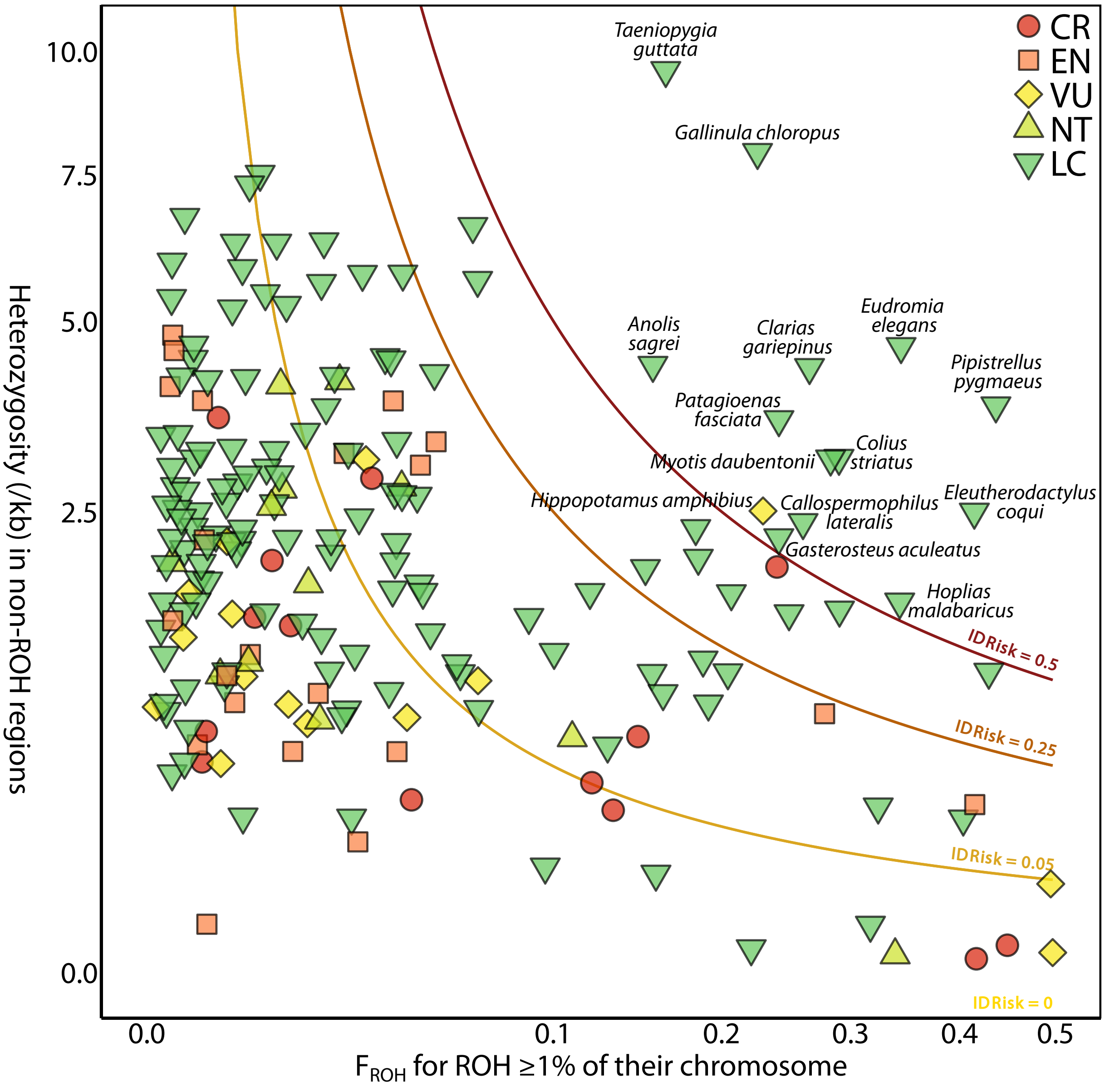


Figure S15: IDRisk values for all species, color coded by risk level. Clines show the boundaries between the different risk categories. Low risk is categorised as below 0.05. Moderate risk is between 0.05 and 0.25. High risk is from 0.25 to 0.5. Extreme risk is anything with an IDRisk value of above 0.5. All species with extreme risk are labelled. LC, Least Concern; NT, Near Threatened; VU, Vulnerable; EN, Endangered; CR, Critically Endangered.
